# A nondegenerate convective-reaction-density-dependent diffusion model of in vitro glioblastoma tumor growth

**DOI:** 10.1101/2020.03.12.988832

**Authors:** Tracy L. Stepien, Erica M. Rutter, Yang Kuang

**Affiliations:** Department of Mathematics, University of Florida, Gainesville, FL, USA; Applied Mathematics Department, University of California, Merced, CA, USA; School of Mathematical and Statistical Sciences, Arizona State University, Tempe, AZ, USA

## Abstract

We propose a model for in vitro glioblastoma multiforme brain tumor growth, which uses density-dependent diffusion to capture both the proliferative and migratory behavior of the cancer cells. The model is compared to well-known experimental data and is analyzed for the existence of traveling wave solutions.

## I. Introduction

Glioblastoma multiforme is a deadly form of brain cancer with a very short mean survival time from detection – less than 15 months (Norden and Wen [4]). Glioblastoma cancer cells exhibit excessive amounts of proliferation as well as migration. It is difficult to effectively treat these tumors, as surgical resection is able to remove the core of the tumor but not the migratory cells. This results in a challenging task to mathematically model all aspects of glioblastoma growth.

Many models of glioblastoma growth include reaction-diffusion equations which accurately capture the proliferating core, such as the early efforts by Tracqui et al. [9] and Swanson et al. [8]. (See also the review paper by Martirosyan et al. [3] and references therein.) These models have also been modified to include spatially dependent diffusion to model more heavily the migratory behavior of the tumor.

Another approach for modeling glioblastoma growth is to separate the glioblastoma cells into two separate populations: the proliferating core cells and the migratory cells, such as in Stein et al. [6]. Their model was based off of observations from in vitro glioblastoma tumor spheroid spreading. In addition to the standard reaction-diffusion terms in the equation for migratory cells, the model includes a radially biased motility term corresponding to convection to account for the situation where cells detect the location of the tumor core and actively move away from it.

We derived a single equation model in Stepien et al. [7] that captures both the migratory and core tumor characteristics as accurately as the dual-equation approach. We analyzed the existence of traveling wave solutions and corroborated the minimum wave speed with simulations. We performed a sensitivity analysis on the parameters in the model to detect how variations in parameters affect the numerical simulation and optimized the parameters according to the experimental data of Stein et al. [6] to validate the model.

## II. Results

### A. Traveling wave speed analysis

A traveling wave solution of (2)–(3) is a solution of the form *u*(*x,t*) = *w*(*x*−*kt*), where *k* ≥0 is the speed of the traveling wave and the function *w*(*z*) is defined on the interval (−∞,∞) and satisfies the boundary conditions w→1 as z→−∞ and w→0 as z→∞. In Stepien et al. [7], we showed that a solution to the boundary value problem that results from substituting the traveling wave ansatz exists via phase plane analysis. This solution exists as long as the speed of the traveling wave is greater than or equal to the minimum wave speed

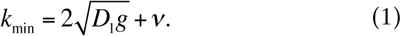

Simulations showed that this minimum wave speed is the observed speed of the traveling wave in some cases, and in other cases the observed speed is faster than the minimum wave speed *k*_min_ (Stepien et al. [7]). In particular, when *D*_2_=0, we found that the observed speed equals the minimum wave speed *k*_min_, but when *D*_2_≠0, the observed speed appears to depend on other parameters besides *D*_1_, *g*, and *ν*. The nonlinear diffusion can be considered as contributing convection with a “velocity” –*D*’(*u*)∂*u*/∂*x*, but the minimum wave speed *k*_min_ is calculated after linearizing which could be the reason for the differences between the speeds.

### B. Parameter Sensitivity and Estimation

To show that the proposed model is viable, we compared numerical simulations with experimental data. The numerical simulations were run over a large spatial domain and boundary conditions specify that there are no tumor cells at the boundaries, in other words, *u*(*x,t*)=0 when *x*=±1 cm. The initial tumor radius is 210 µm and the maximum cell density *u*_max_=4.2×10^8^ cells/cm^3^ (Stein et al. [6]). For the initial condition, we assume that the cell density is 95% of *u*_max_ for the initial core tumor radius of 210 µm and zero elsewhere. Details of the numerical method can be found in Stepien et al. [7].

The base parameters chosen for the parameter sensitivity analysis were *D*_1_=10^−4^ cm^2^/day, *D*_2_=9.99×10^−5^ cm^2^/day, *a*=0.1 cells/cm^2^, *n*=1, *g*=0.5/day, and *ν*=0.01cm/day. To test the sensitivity of one parameter, all the other parameters were held constant and the parameter in question was varied over a physiologically relevant range. The results of this analysis are shown in Fig. 1, and it is apparent that while some parameters are more sensitive than others (*D*_1_, *g*, and *ν*), all parameters are sensitive and we must take care when we estimate optimal parameters.

**Figure 1.**
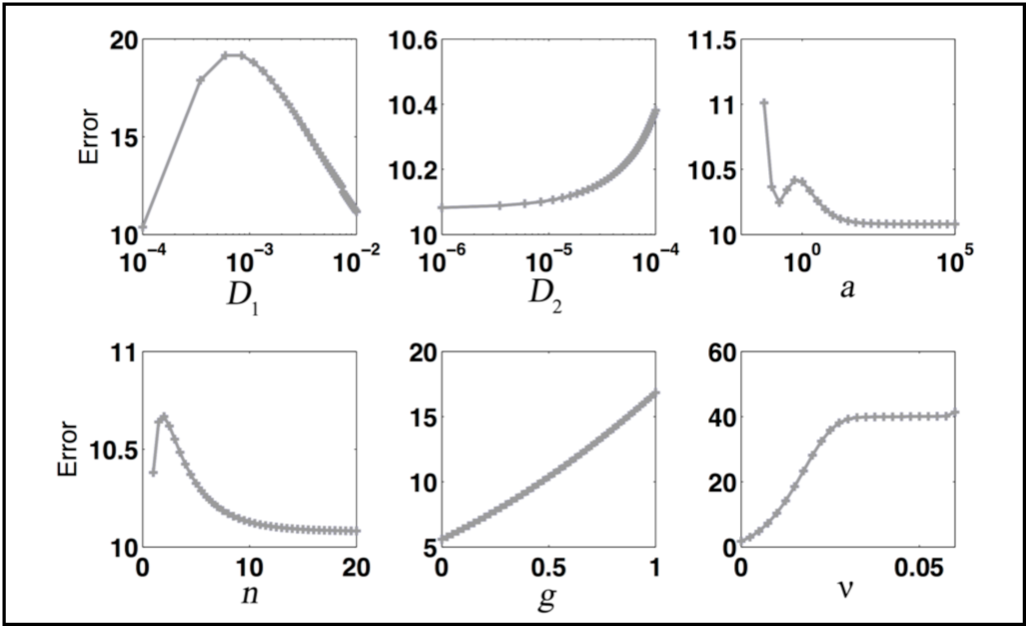
Parameter sensitivity analysis showing the error calculated when comparing the experimental data and numerical simlations.

To estimate optimal parameters that result in numerical simulations that best match the experimental data, we used the MATLAB program `fminsearch`. Various initial parameter guesses were used as input to ensure parameter values were optimal, and the optimized parameters found are given in Table 1.

**Table I.** Table I. Optimized Parameter Values

| Parameter | Optimized Value | Units |
| --- | --- | --- |
| $D_1$ | $5.5408 \times 10^{-6}$ | $\text{cm}^2/\text{day}$ |
| $D_2$ | $5.3910 \times 10^{-6}$ | $\text{cm}^2/\text{day}$ |
| $a$ | 0.021188 | $\text{cells}/\text{cm}^3$ |
| $n$ | 1.2848 | dimensionless |
| $g$ | 0.49120 | $/\text{day}$ |
| $v$ | $4.6801 \times 10^{-5}$ | $\text{cm}/\text{day}$ |

Fig. 2 shows the numerical simulation with parameters from Table 1 compared to the experimental data and model of Stein et al. [6]. Our model is successful in capturing the behavior of both the tumor core cells and the migratory cells and is more accurate than the model of Stein et al. [6]; in fact, our total error is approximately one-half that of the model of Stein et al. [6].

**Figure 2.**
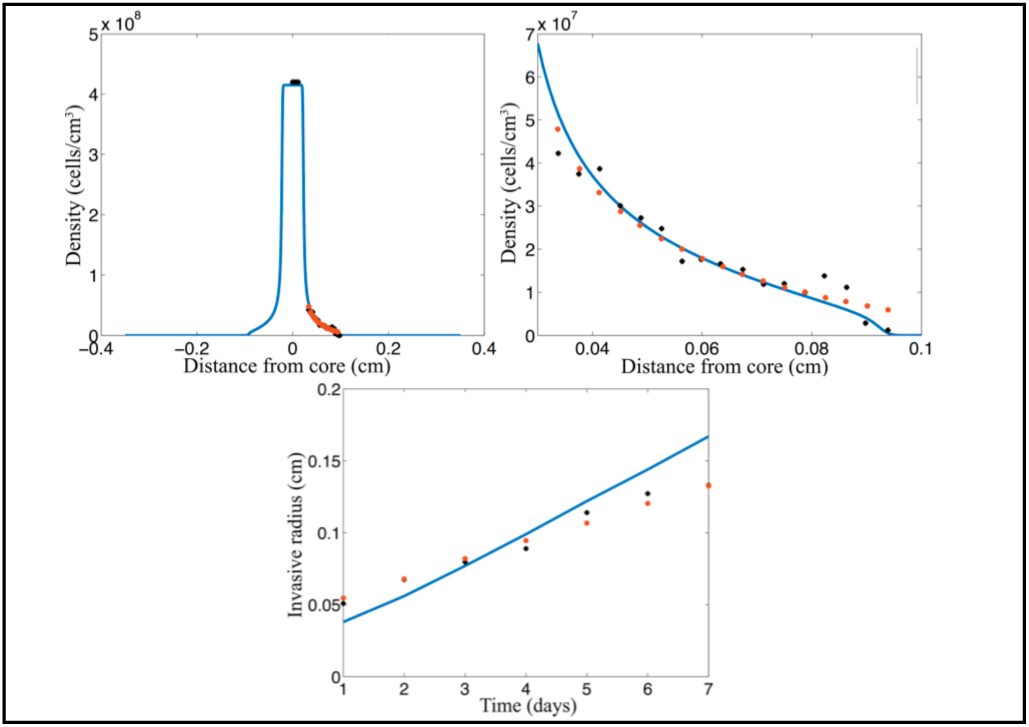
Numerical simulation of the density-dependent diffusion model (2)–(3) and optimized parameters as given in Table 1 (blue) compared to experimental data from Stein et al. [6] (black) and their numerical simluations (red).

## III. Quick Guide to the Methods

The in vitro experiments of Stein et al. [6] involved implanting two human astrocytoma U87 cell lines into gels— one with a wild-type receptor (EGFRwt) and one with an overexpression of the epidermal growth factor gene (ΔEGFR). The tumor spheroids were left to grow over 7 days and imaged every day. In the same study, Stein et al. [6] also developed a mathematical model that assumes the tumor cells leave the tumor core and become invasive cells to invade the collagen gel. The mathematical model that we developed in Stepien et al. [7] is based off of this model of Stein et al. [6].

### A. Equations

In Stepien et al. [7], we consider a density-dependent convective-reaction-diffusion equation,

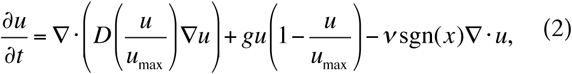

where *u*(*x,t*) is the density of tumor cells, *u*_max_ is the carrying capacity, *g* is the growth rate, *ν* is the degree at which cells migrate away from the tumor core, and *D*(*u*\*) is the density-dependent diffusion function where *u*\*=*u*/*u*_max_. The experimental work from Stein et al. [6] suggests that the diffusion is large for areas where the cell density is small (the migrating tumor cells), but diffusion is small where the cell density is large (the proliferating tumor cells). Thus, to capture this behavior we set

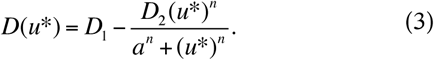

For biologically relevant parameters we assume that *D*_1_, *D*_2_, *g, a*, and *ν* are all positive, *n*>1, and *D*_2_≤*D*_1_ to avoid “negative” diffusion, which is a problem both biologically and numerically. The parameter *n* governs how steeply the density-dependent diffusion function decreases and the parameter *a* governs the *u*\* value at which the transition is occurring at half maximal rate. *D*_1_ and *D*_2_ govern the range of the function.

### B. Type of settings in which these methods are useful

We used this mathematical model to study in vitro glioblastoma growth, but future studies could be used to study in vivo data. The model would then need to include brain geometry. This model could be extended to include more complex behavior such as tumor cell necrosis, brain tissue type differentiation, and mass effect. Instead of density-dependent diffusion, it may be appropriate to implement anisotropy through diffusion tensor imaging (Jbabdi et al. [2], Bondiau et al. [1], Painter and Hillen [5]) or to consider a non-local diffusion with proliferating and dispersing cell groups in which dispersing cells convert proliferating cells into dispersing ones.

